# Maternal low protein diet alters the development of reward circuits from childhood to adulthood by reshaping its function

**DOI:** 10.1101/2025.02.28.639479

**Authors:** Julie Paradis, Chloé Guillaume, Morgane Frapin, Dimitri Meistermann, Anthony Pagniez, Pierre de Coppet, Valérie Amarger, Patricia Parnet, Vincent Paillé

## Abstract

Inadequate nutrition during pregnancy can lead to intrauterine growth retardation and low birth weight, which in turn increases the risk of developing metabolic disorders in adulthood, according to various epidemiological and clinical studies. The inclination of individuals born with low birth weight towards palatable foods indicates a possible modification in the hedonic aspect of their eating behavior. However, our understanding of the ontogenesis of structural organization and function within the brain’s reward circuits remains limited. Therefore, the objective of this research is to investigate the preferences for palatable food, molecular signatures of reward circuits, and functional properties of the nucleus accumbens (NAc) in a rat model of perinatal protein restriction (LP). Starting from weaning, continuing into adolescence and adulthood, a longitudinal analysis was conducted on rats born to mothers with protein-restricted diets during gestation and lactation (LP pups), comparing them to pups born from control dams (CD pups). The LP group exhibited an increased preference for palatable food at day 25 after birth (P25), followed by a decreased preference during adolescence (P50), and no significant difference in palatable food preference at P95 (young adult) compared to CD rats. Molecular and electrophysiological assessments of medium spiny neurons (MSN) in the NAc revealed a reorganization of reward circuits during crucial developmental periods, potentially influencing the attractiveness of palatable food for the LP group. This study represents the first exploration of how preferences for palatable food evolve throughout an individual’s lifespan and how these observations correlate with the remodeling of reward circuits. By shedding light on the molecular and functional aspects of reward circuits, we contribute to a better understanding of the link between perinatal nutrition, behavioral preferences, and the underlying neural mechanisms.

## 1. Introduction

The perinatal period is a critical developmental window in which the growth and development of the fetus and young child are totally dependent on the nutritional, hormonal and metabolic environment provided by the mother. An unbalanced diet during the perinatal period, also referred for Human as the first 1000 days, may lead to metabolic adaptations of the fetus. The unbalanced perinatal diet can be due to overeating or maternal undernutrition and unexpectedly will end up with the programming of higher risk to cardiovascular disorders, diabetes or obesity (Barker 1992; Hales and Baker, 2001; McCance et al., 1994). The concept of Developmental Origin of Health and adult Disease (DOHaD) has been documented for individuals born with Intra Uterine Growth Retardation (IUGR) (Baker, 1992; Green et al., 2010; Lussana et al., 2008; Ravelli et al., 1999) induced by a reduction in nutrient transfer to the fetus (Graccioli and Lager, 2016; Lager and Powell, 2012) but, eventually, also for individuals born to overweight mothers.

Studies in humans and animal models have shown a risk of developing abnormal eating behaviors, in particular changes in food preferences and intake, with an increased preference for fatty and sweet foods at different ages among individuals born with IUGR (Ayres et al. 2010; Barbieri et al., 2010; Bellinger, 2004; Coupe et al., 2009, 2012; Dalle Molle et al., 2015; Lussana et al., 2008; Martin-Agnoux et al., 2014; Migraine et al., 2013; Oliveira et al., 2015; Portella et al., 2012). However, the cellular and molecular mechanism involved in such preferences are poorly understood. The central reward circuits that control the hedonic and motivational aspect of food intake appear to be a good candidate that could be impacted by nutritional programming as the dopaminergic system is altered in obese individuals in both humans and animals (Alsïo et al., 2010; Davis et al., 2008; Haltia et al., 2007; Stice et al., 2008) and the plasticity of these circuits is altered in animal model that have been exposed to a high-calorie diet during adolescence (Naneix et al., 2016; Vendresculo et al., 2010). Indeed, the mesocorticolimbic dopaminergic circuit matures late and its development extends from the end of fetal life, childhood with dietary diversification and even until adolescence influenced by more personalized food choices. (Zestler, 2018). This period of intense plasticity makes this circuit sensitive to remodeling due to maternal perinatal feeding or postnatal diet. A modification of the expression of DA receptor, or of the enzymes involved in DA synthesis, could be demonstrated in rodent model of IUGR (Alves et al., 2015; Dalle Molle et al., 2015; Laureano et al., 2016, 2018; de Melo-Martimiano et al., 2015; Vucetic et al., 2010b). Most of these studies are performed in adult animals and therefore described the consequences of perinatal diet on adult reward circuits. Little is known on the ontogeny of the dopaminergic circuit following perinatal undernutrition and how early modifications of this circuit orient food preferences during development. We aimed at understanding how eating behavior evolves during the 3 critical phases of postnatal development (childhood P25, adolescence P50 and adult P100) following perinatal undernutrition. We previously conducted some work on the consequences of perinatal maternal overfeeding on the development of the dopaminergic and the GABAergic system of rat pups. We showed in the offsprings a clear evolution for fat preference and that maternal western diet intake has a long-lasting influence on the organization of the homeostatic and hedonic circuits regulating eating behavior (Paradis et al., 2017; Lippert et al., 2020).

The hedonic and/or motivational component of eating behavior involves GABAergic neurons from brain regions responsible for addictions such as the nucleus accumbens (NAc) since the NAc is mainly composed of GABAergic MSN neurons that will integrate mesocorticolimbic system information, dopaminergic afferences of VTA and cortex glutamatergic afferences, to convert them into action via projections to the ventral pallidum and other structures (Hu et al., 2016; Salgado and Kaplitt, 2014). However, the electrophysiological properties of MSN neurons and the communication within these reward circuits have not been documented in the case of perinatal undernutrition, nor are their temporal evolution. Therefore, we analyzed the influence of maternal protein restriction on (i) palatable food preferences, (ii) gene expression of the dopaminergic and gabergic systems, (iii) electrophysiological properties of GABAergic neurons in the reward system on rat offspring from birth to adulthood. For this purpose, a longitudinal study (from childhood P25, through adolescence P45 to the young adult P95 stage) was conducted in rats born IUGR following a maternal protein restriction. Palatable food preference tests were performed, followed by molecular analysis (RNA-seq, qPCR) of the different markers involved in the systems regulating food choice, motivation and intake, and correlated with electrophysiological analysis (patch-clamp) of MSN neurons in NAc, a key structure involved in preference and motivation. Our results highlight the reorganization of reward circuits at three key stages in an individual’s life following nutritional programming during the perinatal period.

## 2. Results

### 2.1. Body Weight and Growth

Maternal protein restriction during gestation (from G1 to G21) significantly modified pups body weight at birth (CD: 6.905 ± 0.045 g vs LP: 6.205 ± 0.086 g p<0.0001) (Figure 2 B). Body weight gain from birth to weaning was 58.9% lower in offspring from LP dams with a body weight significantly lower at weaning in offspring born from LP dams (CD: 42.76 ± 0.860 g vs LP: 27.34 ± 0.670 g p<0.0001) (Figure 2 A). From weaning to the end of the experiment (P95), the rats were fed with standard chow diet and body weight remained lower for LP group. See details on Figure 2: Childhood (P25) CD: 45.64 ± 1.333 g vs LP: 38.77 ± 1.424 g p= 0.0009, Adolescence (P50) CD: 177.7 ± 5.089 g vs LP: 160.1 ± 6.072 g p= 0.0297, and at Younf adult (P95) CD: 443.3 ± 11.33 g vs LP: 373.8 ± 14.24 g p= 0.0006 (Figure 2 A, C, D and E).

**Figure 1.**
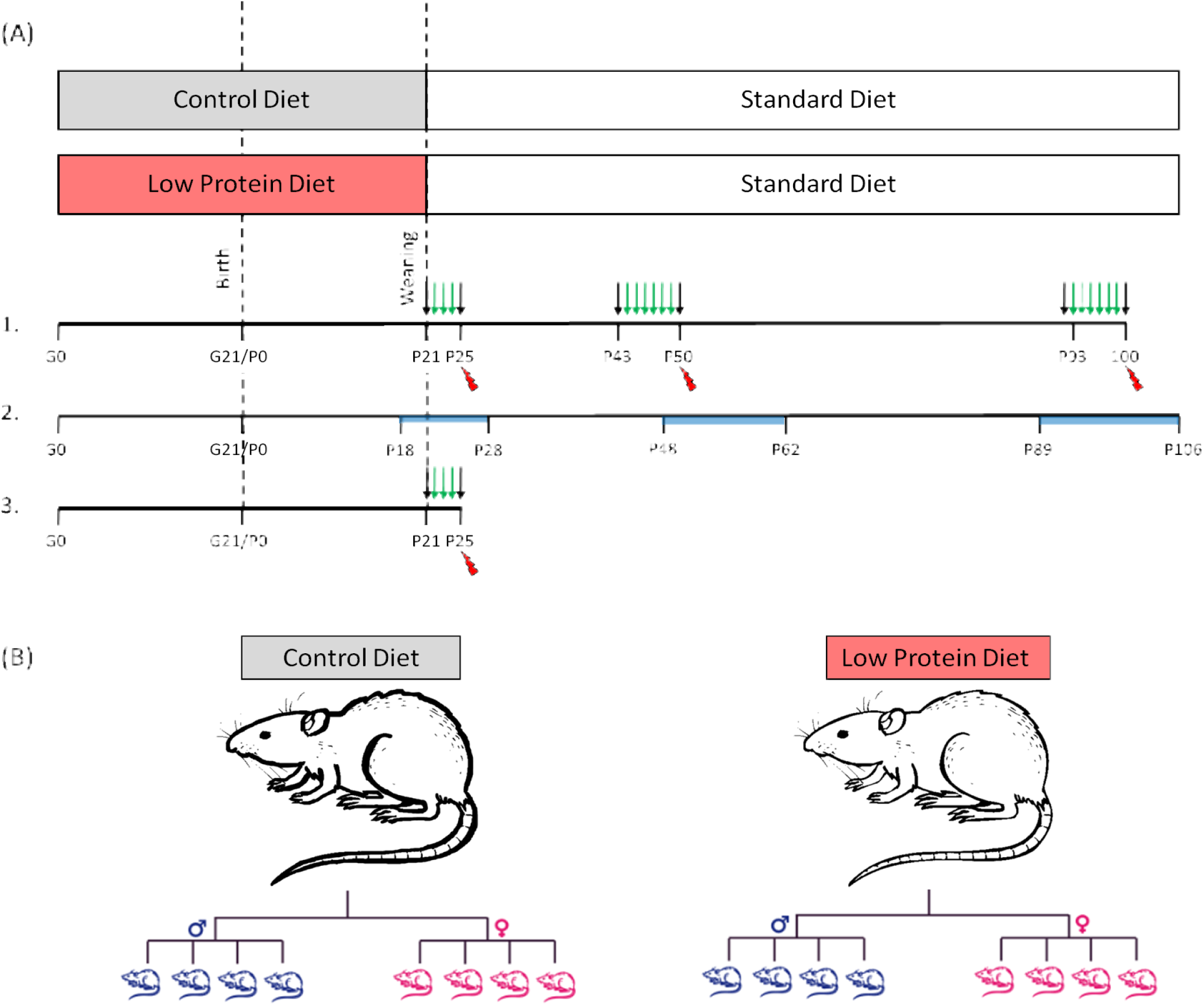
Experimental design **(A)** Schematic overview of the study. On gestational day 1 (G1), female Sprague-Dawley (SPD) rats were assigned to either a control diet or a low-protein (LP) diet throughout gestation and lactation. At weaning, all offspring were placed on a standard chow diet until the end of the experiment. Three main study phases were conducted: 1. At three developmental stages—P25 (childhood), P45 (adolescence), and P95 (young adulthood)—a fat preference test was performed for 1 day (P25) or 3 days (P45 and P95) (N = 12 per group). Black arrows indicate a habituation period with free access to a normal diet in individual cages, and green arrows indicate the 24 h Western Diet (WD) preference test. After this test, rats were sacrificed (red symbol). Half of them (n = 6 per group/age) were perfused with 4% paraformaldehyde, and brains were collected for immunohistochemistry. The other half (n = 6 per group/age) served for plasma measurements and molecular analyses. 2. The same design was used to investigate the electrophysiological properties of nucleus accumbens (NAc) medium spiny neurons (MSNs) at the three time points. 3. D1/D2 neuron quantification protocol: SKF81297 (5.0 mg/kg) or Raclopride (0.3 mg/kg) was administered by intraperitoneal injection on Day 4 after a free-choice test with high-fat food in P25 pups (as described above). **(B)** At birth, litter size was standardized to eight pups with a 1:1 male-to-female ratio, and only males were used in this study. For each time point (P25, P45, P95), one male per litter was used for behavioral testing and subsequent postmortem analyses.

**Figure 2.**
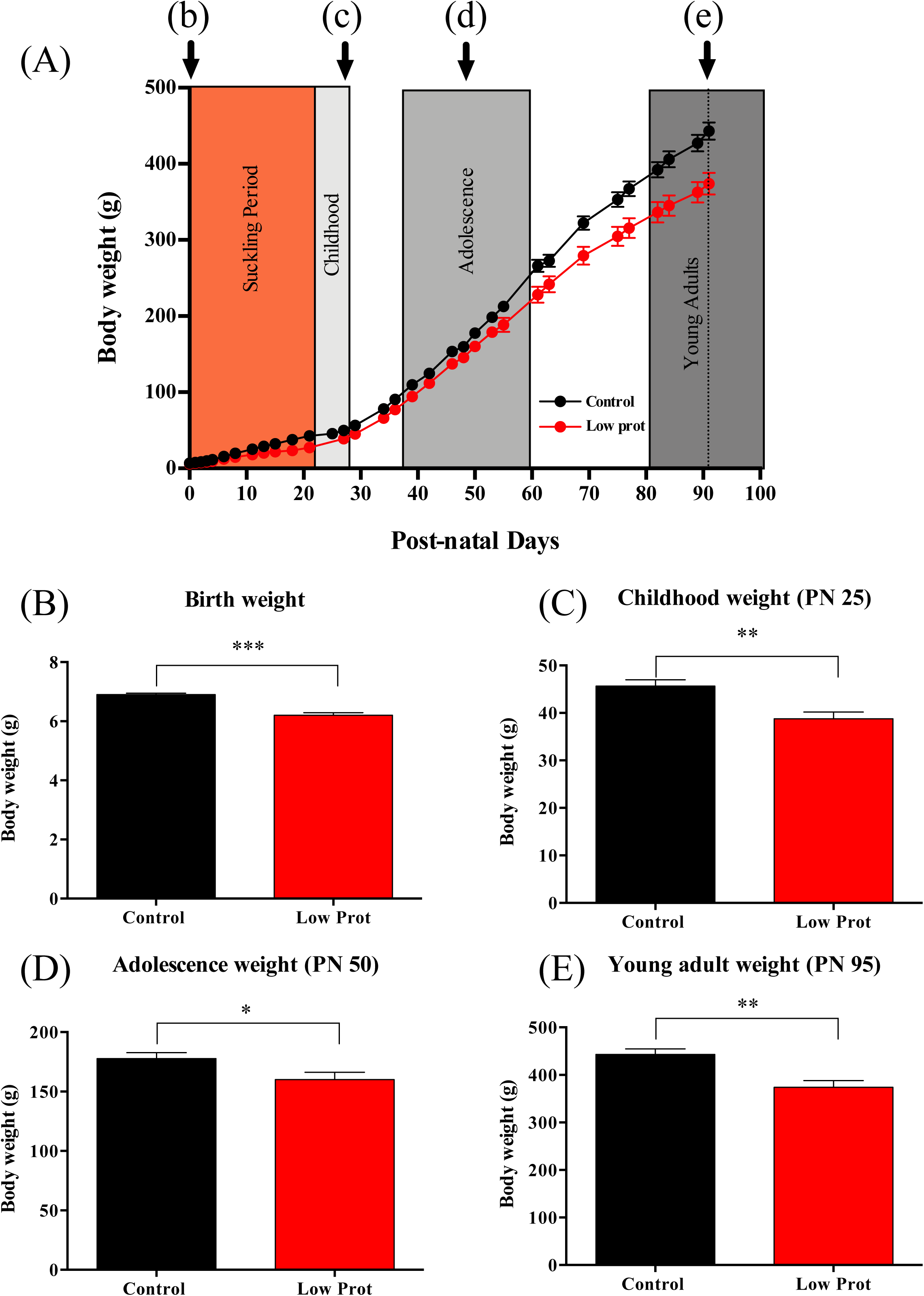
Body weight. **(A)** Average body weight across the three post-weaning periods. **(B)** Birth weight (Mann–Whitney test, p < 0.0001; Control n = 40, IUGR n = 44). **(C)** Childhood weight (P25) (Mann–Whitney test, p = 0.0017; Control n = 30, IUGR n = 25). **(D)** Adolescent weight (P50) (Mann–Whitney test, p = 0.0405; Control n = 30, IUGR n = 25). **(E)** Young adult weight (P95) (Mann–Whitney test, p = 0.0014; Control n = 18, IUGR n = 12). Data are presented as mean ± SEM.

### 2.2. Impact of Perinatal Protein Restriction on Fat Preference from Weaning to Adulthood

To explore the impact of the perinatal low-protein diet on preference for the Western diet (WD), we used a free food choice paradigm at three different time points during development. This test was used to specifically investigate preference for a palatable food (fat and sugar or western diet).

Regarding preference for WD, we showed a significantly higher preference for WD on the second and third test day in the LP group at P25 (day 2 CD: 90.21±3.541% vs. LP: 93.38±6.304% p=0.0477; day 3 CD: 86.77±6.140% vs. LP: 98.87±0.892 p=0.009). At P50, the effect was completely reversed, LP rats had a lower preference for WD on day 4, 5 and 6 of the test compared to the CD group (day 4 CD: 91.21±4.958% vs LP: 72.66±7.894% p=0.013; day 5 CD: 91.69±3.079% vs LP: 68.79±7.152% p=0.006; day 6 CD: 91.14±3.439% vs LP: 70.36±4.267% p=0.001). Finally, at P100, a smaller difference for WD was found only day 1 for the LP group (day 1 CD: 86.22±3.994% vs. LP: 96.61±1.435% p=0.0277) but no further differences were observed for the remaining five days of the test (Figure 3 A).

**Figure 3.**
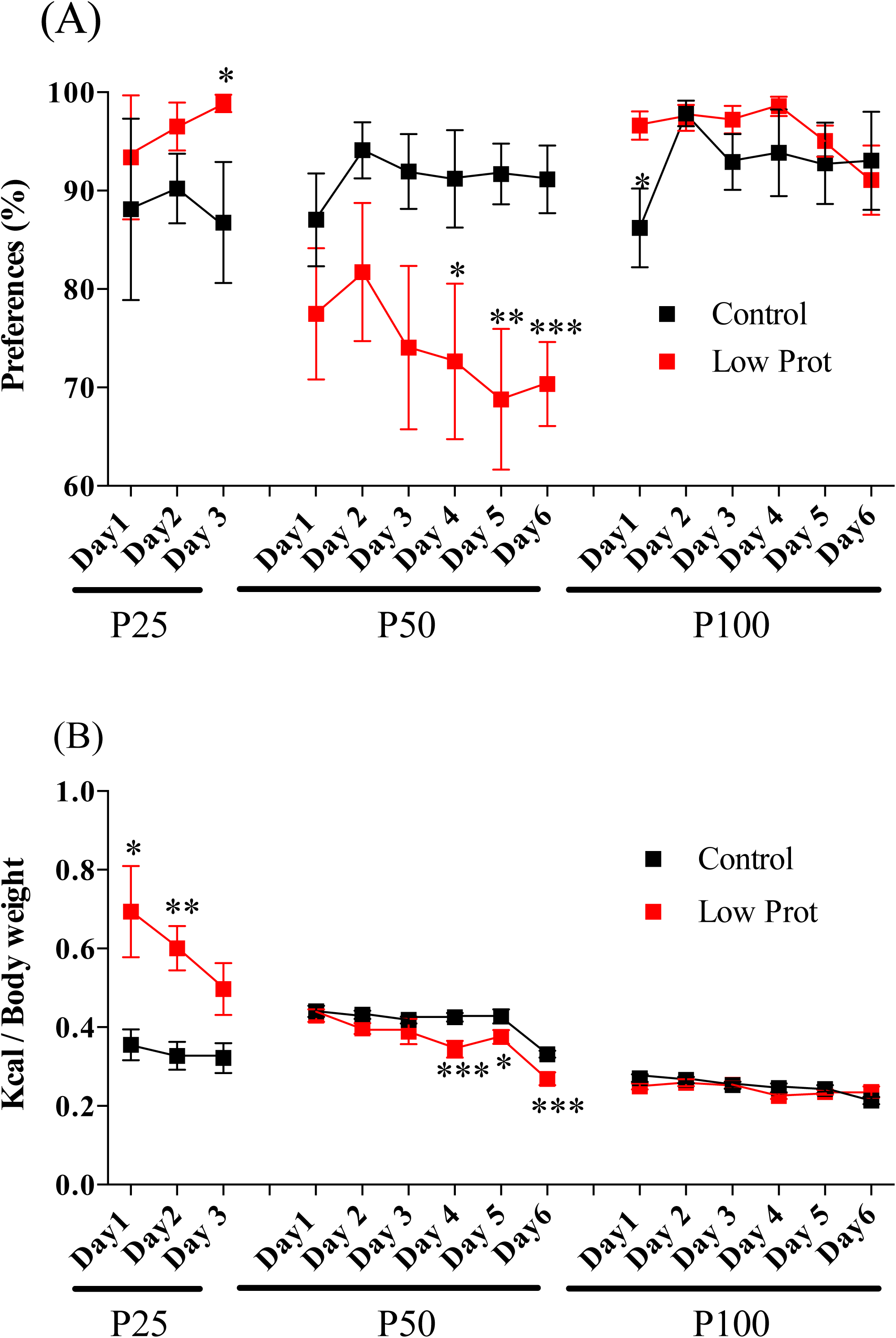
Free-choice paradigm with ad libitum Western Diet (WD) vs. Standard Diet (SD) **(A)** Consecutive days of fat preference testing on the same set of animals at P25, P50, and P100. **(B)** Calorie intake during consecutive days of fat preference testing at P25, P50, and P100. Data are shown as mean ± SEM. Different animal cohorts were used at each age (n = 6 per group). Mann–Whitney test: *p < 0.05, **p < 0.01, ***p < 0.001.

Concerning the total energy intake during the test, the results confirm those obtained during the preference analysis. Indeed, we observed a significantly higher consumption of total Kcal on the first and second day of the test in the LP group at P25 (day 1 CD: 0.355±0.039Kcal/g body weight vs. LP: 0.694±0.115 Kcal/g body weight p=0.022; day 2 CD: 0.327±0.035Kcal/g body weight vs. LP: 0.601±0.056 Kcal/g body weight p=0.002). At P50, LP rats had significantly lower total energy intake on the fourth, fifth and sixth days of the test compared to the CD group (day 4 CD: 0.425±0.011 Kcal/g body weight vs. LP: 0.344±0.021 Kcal/g body weight p=0.0007; day 5 CD: 0.427±0.017Kcal/g body weight vs LP: 0.375±0.017 Kcal/g body weight p=0.044; day 6 CD: 0.331±0.008Kcal/g body weight vs LP: 0.269±0.016 Kcal/g body weight p=0.0007). No difference in total energy intake was found at P100 (Figure 3 B). We decided then to explore, only in males, molecular and physiological modifications of the reward circuits.

### 2.3. Impact of Perinatal LP Diet on Electrophysiology Properties of NAc MSN on males

#### 2.3.1. Action Potential recording

Different parameters were analyzed (Table 2) reflecting passive properties of neurons (input resistance, resting membrane potential, capacitance) and active properties of MSNs (rheobase, spike characteristics, spike frequency). Among passive properties, no differences were found between LP and CD rat MSNs during childhood and adolescence (see Table 2). A trend toward lower input resistance was observed in LP rat MSNs (CD: 163. ± 12.40 mOhm vs. LP: 138.2 ± 15.29 mOhm p=0.0589) (see Table 2). Among the active properties, the action potential threshold was significantly higher in LP rat MSNs (CD: −39.65 ± 0.525 mV vs. LP: −37.05 ± 0.501 mV p=0.0018) and the duration of sAHP was longer in the LP group (CD: 60.84 ± 4.644 ms vs. LP: 73.99 ± 5.285 ms p=0.0266) during childhood (see Table 2). In adolescence, the spike rise/decay time ratio was significantly higher in the LP group (CD: 0.330 ± 0.064 vs. LP: 0.329 ± 0.016 p=0.0489) (see Table 2) with a trend toward lower total spike duration and lower spike decay time in MSNs of LP rats (results near statistical significance with CD: 3. 169 ± 0.183 ms vs. LP: 2.745 ± 0.097 ms p=0.0715 and CD: 2.438 ± 0.163 ms vs. LP: 2.078 ± 0.088 ms p=0.0665 respectively) (see Table 2). In adulthood, rheobase is significantly higher in MSNs from LP rats (CD: 100.8 ± 6.088 pA vs. LP: 141.2 ± 13.39 pA p=0.0068) and both sAHP duration and spike rise time were significantly lower in the MSNs from LP group (CD: 80.64 ± 4.997 ms vs. LP: 67.32 ± 3.555 ms p=0.0475 and CD: 0.8606 ± 0.046 ms vs. LP: 0.6831 ± 0.041 p=0.0059 respectively) (see Table 2). Furthermore, the I/V curves were similar between LP and CD rat MSNs during childhood and adolescence but statistically different in adulthood with a more pronounced inward rectification in LP rat MSNs and a smaller I/V curve slope beyond 0pA in LP rat MSNs (Figure 4 D, E, F).

**Figure 4.**
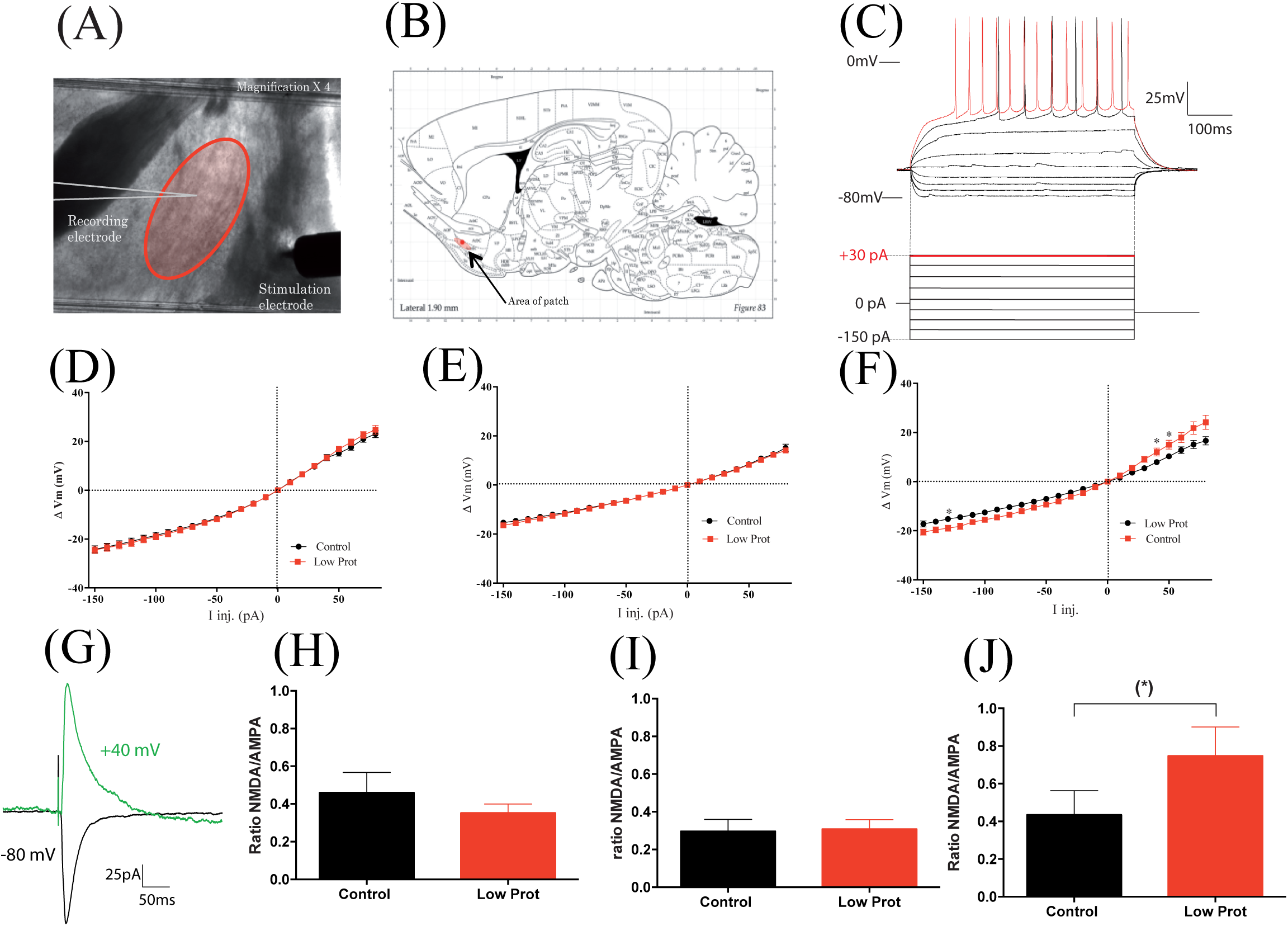
Electrophysiological properties. **(A)** Photomicrograph of the patch area. **(B)** Sagittal view adapted from Paxinos and Watson showing the recorded region. **(C)** Representative medium spiny neuron (MSN) spiking pattern in current clamp (red trace = rheobase + 30 pA). **(D, E, F)** Current-voltage (I–V) curves for MSNs at P25 (Ctrl n = 30, LP n = 26), P50 (Ctrl n = 16, LP n = 20), and P100 (Ctrl n = 12, LP n = 17). **(G)** Representative excitatory postsynaptic current (EPSC) traces at −80 mV (black) and +40 mV (green). **(H, I, J)** NMDA/AMPA ratio at P25 (Ctrl n = 15, LP n = 13), P50 (Ctrl n = 11, LP n = 12), and P100 (Ctrl n = 9, LP n = 13).

#### 2.3.2. NMDA/AMPA ratio

To determine whether protein intake deficiency during gestation and lactation impacted MSNs plasticity, the NMDA/AMPA ratio was measured as previously described (Dumont et al. 2005). For this purpose, EPSCs were recorded in the NAc MSNs obtained by cortical stimulation of glutamatergic presynaptic fibers (cf methods). No differences were found in NMDAR/AMPAR ratios at P25 and P50. A tendency for a higher NMDAR/AMPAR ratio was observed at P100 (CD: 0.434 ±0.127 vs LP: 0.748 ±0.153 p=0.059) (Figure 4 G-J).

#### 2.3.3. Miniature spontaneous GABAergic current recording

We decided to evaluate the impact of perinatal undernutrition on GABAergic signaling in MSNs at the different critical stages of our study. As previously described, there are 2 types of ionotropic GABAergic signaling that differ according to GABAAR localization: tonic signaling (synaptic localization) and phasic signaling (extrasynaptic localization) (Kaneda et al., 1995; Farrant and Nusser, 2005; Glykys and Mody, 2007; Brickley and Mody, 2012). These 2 types of current have a different impact on neuronal excitability (Farrant and Nusser, 2005; Glykys and Mody, 2007). To identify these currents, we used an intracellular cesium-based solution (see Methods) associated with the progressive application, in the perfusion bath, of ionotropic glutamatergic receptor antagonists (50 mMD-AP5 and 10 mMCNQX) to which we added a specific GABAARs inhibitor (100µM PTX) (Figure 5A). This last drug (PTX) thus allowed us to identify the tonic GABAergic component (see method).

**Figure 5.**
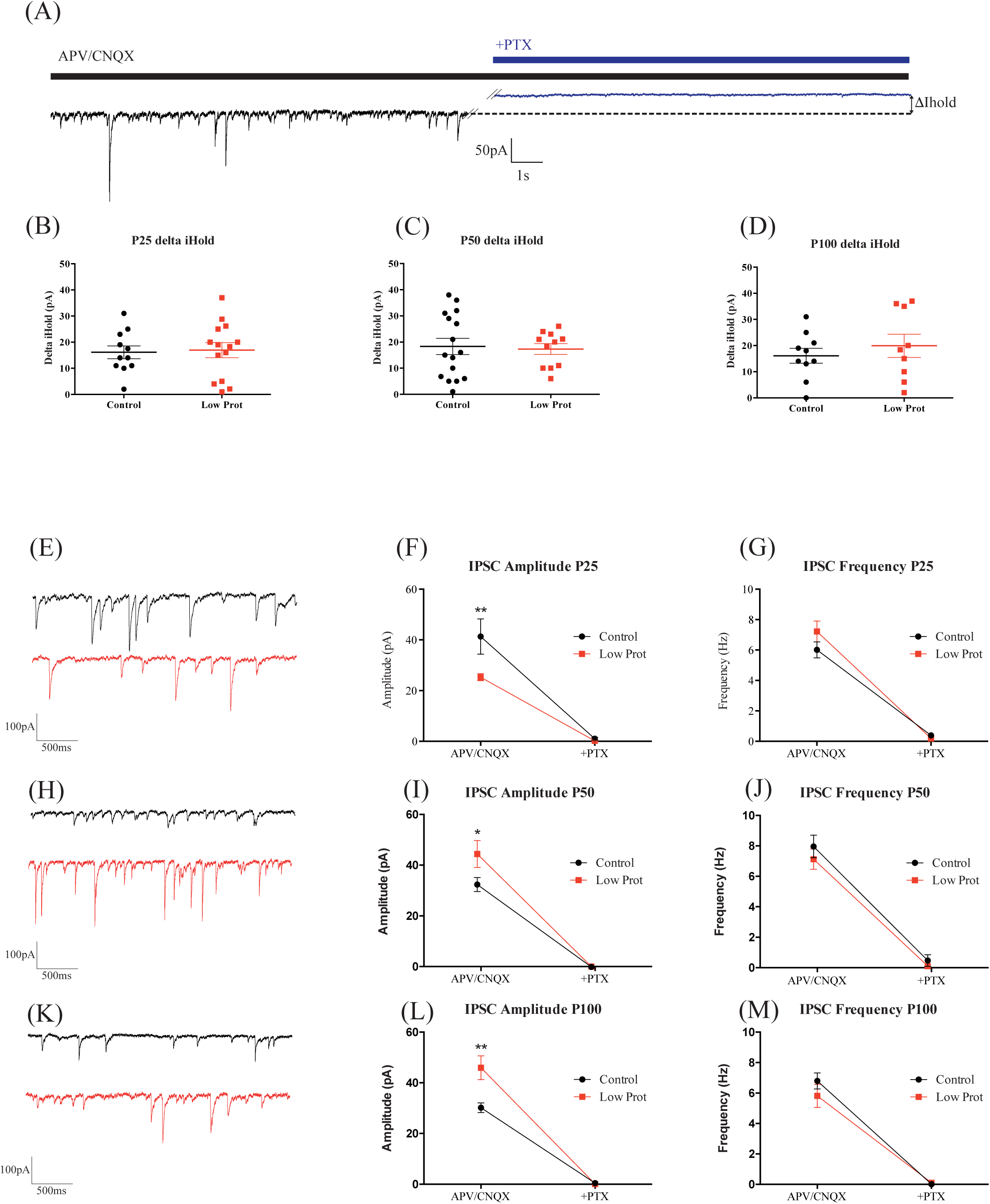
GABAergic currents. **(A)** Raw trace recordings of MSNs in the presence or absence of picrotoxin (in D-AP5/CNQX). Picrotoxin blocks both tonic and phasic GABA_A receptor-mediated currents. **(B, C, D)** Tonic GABA_A current determined by the difference in holding current (ΔI_hold) before and after picrotoxin at P25 (Ctrl n = 11, LP n = 14), P50 (Ctrl n = 16, LP n = 11), and P100 (Ctrl n = 10, LP n = 9). **(E–G), (H–J), (K–M)** Representative traces of miniature inhibitory postsynaptic currents (mIPSCs) measuring GABAergic currents (frequency and amplitude) at P25 (Ctrl n = 11, LP n = 14), P50 (Ctrl n = 16, LP n = 11), and P100 (Ctrl n = 10, LP n = 9). Data are shown as mean ± SEM. Mann–Whitney test: *p < 0.05, **p < 0.01, ***p < 0.001, (*p < 0.07).

During infancy (P18-28), the amplitude of the IPSC (phasic current) was significantly lower for LP rat MSNs (CD: 41.33 ± 6.94 pA vs LP: 25.30 ±1.38 pA p=0.007) (Figure 5 E, F). In contrast to childhood, the amplitude of IPSCs was higher for LP rat MSNSs in adolescence(P48-62) (CD: 32.73 ± 2.77 pA vs. LP: 44.79 ± 5.29 pA p=0.051) (Figure 5 H, I) and adulthood (P89-106) (CD: 30.18 ± 1.88 pA vs. LP: 45.91 ± 4.70 pA p=0.008) (Figure 5 K, L). No difference in IPSC frequency was found at the three different stages studied (Figure 5 G, J, M).

The tonic GABAergic current, measured after addition of PTX and the subsequent GABAAR blockade, was identical in the 2 groups (CD and LP) at all 3 ages (Wilcoxon signed rank p<0.05 corresponding to the red asterisks on graphs B to D) and was of the same intensity (delta iHold: P25: CD: 16.14± 2.44 vs LP: 16.96± 2.88); delta iHold P50: CD: 18.3± 3.01 vs LP: 17.32± 2.01); (delta iHold: P100: CD: 16.1± 2.82 vs LP: 19.93± 4.44) (Figure 5 B, C, D).

### 2.4. Molecular Signature of Dopamine Circuits in Reward Pathways

To determine whether maternal LP intake during perinatal period has an impact on the reward pathways of the offspring we performed RNA-seq technic on NAc and VTA structures and searched for general enrichment for several different molecular pathway. Screening was performed on tissues obtained after the free food preference choice test at P25, P50 and P100 on six males CD and six males LP. We observed a strong modulation of epigenetic and adhesion pathways for the different ages in VTA and in NAc for the epigenetic pathway. A strong modulation of the dopaminergic pathway was observed in NAc at P25 but this modulation is less important at P50 and P100. Moreover, a strong modulation of GABAergic pathways was noticed in the NAc at P25 and P50 and in the VTA at P100 (Figure 6 A and B). We used RT-qPCR technic to analyze in more details the abundance of some dopaminergic and GABAA receptors molecular markers in the NAc and the VTA at the three time periods in both group (Figure 6 C). At P25, molecular expression of Drd2, Th and DAT were higher in VTA (DrD2 LP/CD difference: 0.077 ± 0.008 p=0.0403; Th LP/CD difference: 0.120 ± 0.0409 p=0.0022; DAT LP/CD difference: 0.079 ± 0.026 p=0.0179), Drd2 and DrD1a molecular expression were higher in NAc of LP offspring (DrD2 LP/CD difference: 0.2130 ± 0.006 p=0.0022; DrD1a LP/CD difference: 0.0734 ± 0.011 p=0.022). At P50, Drd2 and DAT molecular expression were still higher in the VTA (DrD2 LP/CD difference: 0.062 ± 0.016 p=0.0170; DAT LP/CD difference: 0.090 ± 0.027 p=0.0271) but no more differences were observed in the NAc for LP group (Figure 6 B). At P100, Drd2 molecular expression was lower in VTA (DrD2 LP/CD difference: −0.1535 ± 0.058 p=0.0464) and DrD1a expression were higher in the NAc of LP offspring (DrD1a LP/CD difference: 0.051 ± 0.093 p=0.0262) (Figure 6 B).

**Figure 6.**
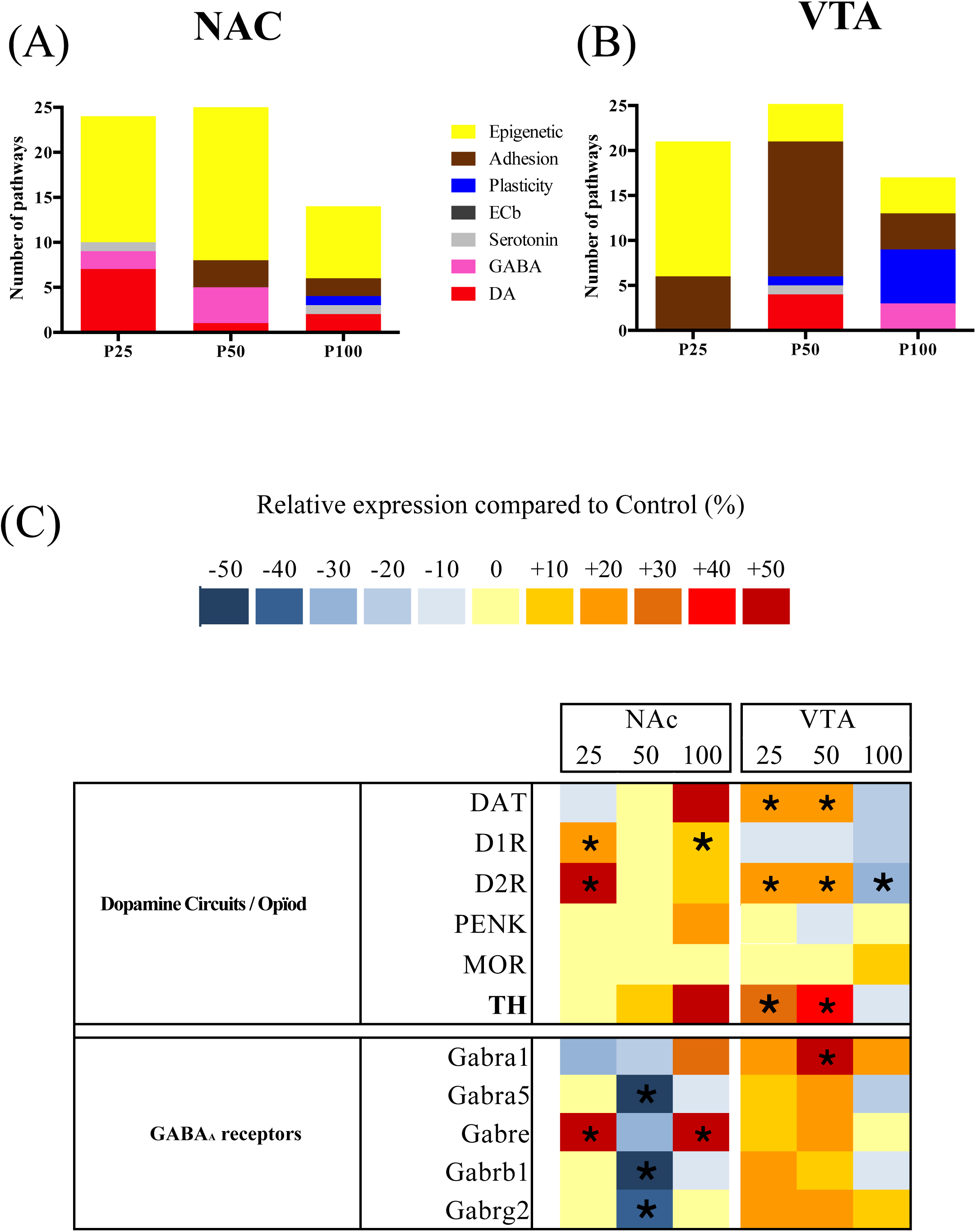
Molecular analysis. **(A)** Number of enriched pathways in different gene families in the ventral tegmental area (VTA) and the nucleus accumbens (NAc) at P25, P50, and P100 (gene family p < 0.01). For each age: CD n = 6, LP n = 6. **(B)** qPCR results (LP expression difference compared to the CD group) for various dopaminergic markers in the VTA and NAc at P25, P50, and P100. Data are presented as mean ± SEM. Mann–Whitney test: *p < 0.05, **p < 0.01, ***p < 0.001, (*p < 0.07).

### 2.5. Quantification of D1R and D2R Neuron Activation in NAc

The NAC MSN neurons are segregated according to the expression of the D1 and D2 receptors corresponding to the direct and indirect pathway respectively (Hersch et al., 1995; Valjent et al., 2009; Gerfen. 2022). In the NAC the segregation is less pronounced than in the rest of the striatum with 20% co-expression of these receptors. Several studies have shown that these neurons have different electrophysiological properties (Surmeier et al., 2007) others indicate that this segregation is not so clear (Bertran-Gonzales et al., 2010). In this study, we were unable to correlate our electrical recordings with the expression of type 1 or 2 dopamine receptors. However, the analysis of our results on the different parameters studied does not show the presence of two populations of neurons with distinct properties. Indeed, the point clouds corresponding to the different parameters are homogeneous. However, we wanted to appreciate the balance between D1 positive and D2 positive neurons. C-Fos positive cell were counted in the Core and shell part of NAc after selective activation of D1 or D2 pathways with administration of agonists (SKF81297 and Raclopride respectively). The results are showed in Figure 7. Activation of D1 or D2 cells is statistically the same in LP rats at PN25 both in the core and the shell (Figure 7, f). In the NAc of control animals, the number of cells activated by the D1 agonist is about 4 times greater in the shell and 3 times greater in the core than the number of cells activated by the D2 agonist (D1/D2 ratio: 3,9 in the Shell and 2,9 in the Core). Perinatal exposure to a low protein food seems to decrease this imbalance to reach the same ratio in the core and shell around 2 times (D1/D2 ratio: 2,2 in the Shell and 2,3 in the Core). (Figure 7).

**Figure 7.**
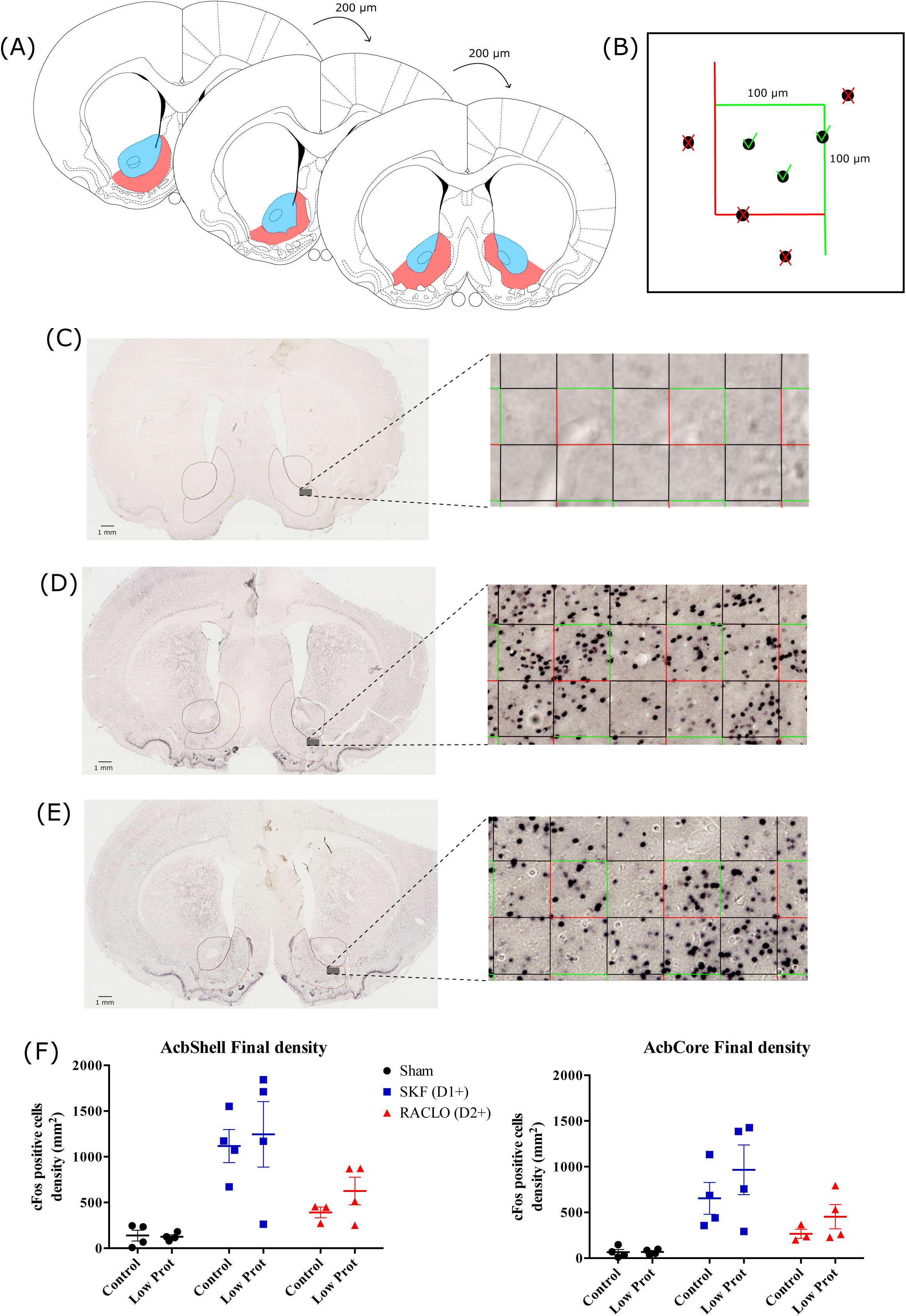
Quantification of dopaminergic D1 and D2 neurons (density and ratio) in the NAc shell and core at P25. (n = 4 per group). **(A)** Slice selection and spacing: based on the Paxinos atlas, slices range from Bregma 1.20 mm to 0.70 mm, with a 200 µm interval between sections. **(B)** Example of the square used for stereological cell counting. Neurons within the inclusion region (inside the square or on the green edges) are counted, while those on or beyond the exclusion edges (red line) are not. **(C)** Representative sham condition: cFos immunohistochemistry (left) and its stereological grid (right). **(D)** Representative raclopride condition: cFos immunohistochemistry (left) and its stereological grid (right). **(E)** Representative SKF81297 condition: cFos immunohistochemistry (left) and its stereological grid (right). **(F)** Density of cFos-positive cells (per mm²) in control or low-protein (LP) perinatal conditions across the three treatment groups (Sham, Raclopride, and SKF).

## 3. Discussion

Our central hypothesis was that a diet restricted in protein during the perinatal period would reshape how the offspring’s reward circuits develop, thereby influencing energy balance, eating behaviors, and food choices. We carried out a detailed study of how maternal low protein (LP) diet exposure during pregnancy and lactation affects dopaminergic (DA) and GABAergic neurons within the ventral tegmental area (VTA) and nucleus accumbens (NAc), from early postnatal life (childhood) through adulthood. Overall, our findings indicate that administering a protein-deficient diet solely in the perinatal window alters early consumption of high-fat foods in the offspring, and this shift is linked to changes in both gene expression and neuronal activity in mesolimbic areas central to reward. Notably, while a pronounced affinity for fatty or sugary foods is observed in childhood, providing a balanced (chow) diet shortly after weaning diminishes the preference for these palatable foods during adolescence, eventually resulting in normalized preferences by adulthood. However, even when dietary preferences no longer deviate from control levels in adulthood, offspring that experienced intrauterine growth restriction (IUGR) exhibit ongoing reorganization in the reward circuitry (particularly in the NAc), persisting beyond the juvenile period.

The most immediate effect of perinatal LP intake was a pronounced IUGR, evidenced by a lower birth weight in pups from LP dams and a roughly 58% decrease in body weight gain by the end of lactation compared with control diet (CD) animals. Prior studies similarly report that protein restriction during lactation triggers extrauterine growth retardation (Coupé et al., 2012; Remmers, 2008; Zhang et al., 2013). After weaning onto a normal diet, we observed no substantial catch-up growth in the LP group, which continued to weigh less than controls throughout the study. These data corroborate earlier reports that various maternal undernutrition models (protein or calorie restriction) tend to produce offspring with reduced birth weights (Breton et al., 2009; Coupé et al., 2012; Da Silva et al., 2016; Orozci-Solis et al., 2009; Zhang et al., 2013). Moreover, our findings align with literature indicating that nutritional restriction spanning the lactation period prolongs low body weight into adulthood (Coupé et al., 2012; Da Silva et al., 2016; Remmers, 2008; Orozco-Solis et al., 2009). Such low weight is potentially attributable to altered milk output or composition by the mother, as described previously (Grigor et al., 1987; Martin-Agnoux et al., 2015).

Interestingly, perinatal malnutrition has been associated with lower attraction to highly palatable foods (Alves et al., 2015; Dalle Molle et al., 2015; Melo-Martimiano et al., 2015). Here, we pursued a longitudinal approach to assess fat preference in offspring raised on a regular diet after weaning.

### 3.1. Impact of Prenatal LP Diet on Childhood (Post-Weaning)

Numerous investigations into perinatal undernutrition models have typically examined hedonic eating behaviors in relatively young adult rodents (Alves et al., 2015; Laureano et al., 2016, 2018; de Melo-Martimiano et al., 2015; Vucetic et al., 2010), thus leaving a gap in knowledge about reward circuitry responsiveness during earlier developmental stages. Given that the brain regions controlling reward are not yet mature at weaning (Gugusheff et al., 2015; Zestler, 2018), it is particularly informative to assess early-life preferences for palatable foods and correlate these behaviors with gene expression profiles in not-yet-fully-developed structures (Gugusheff et al., 2015; Zestler, 2018). Immediately after weaning, LP offspring exhibited a significantly higher inclination toward palatable food intake relative to controls. Similar heightened hedonic responses have been documented in neonates following gestational caloric restriction (Laureano et al., 2016), while elevated fat consumption at weaning has also been observed (Whitaker et al., 2012). Notably, these findings are consistent with clinical observations in children born with IUGR (Ayres et al., 2012; Migraine et al., 2013; Oliveira et al., 2015).

Our results also reveal that, relative to their body weight, LP pups consumed more total calories than controls, suggesting an attempt at post-weaning catch-up. Indeed, LP offspring gained about 41% of their body weight between postnatal day (P) 21 and P25, whereas CD pups only increased by 6% over the same timeframe. Despite this spike in food intake, LP animals still did not achieve the same body weight as controls, supporting the notion that overeating and enhanced preference for rewarding foods may constitute an adaptive mechanism to compensate for deficits incurred during gestation and lactation.

To clarify the underlying molecular changes, we performed differential gene expression RNA-sequencing (DGE-RNA-seq) in the VTA and NAc in 5–6 males per group across three developmental stages. Due to statistical constraints, the power to detect subtle gene-level fluctuations was limited, but gene set enrichment analysis (FGSEA) highlighted dopaminergic and GABAergic pathway alterations in the NAc of LP pups at P25. TAQMAN assays combined with the RNA-seq data pinpointed a marked rise in D2 receptor (D2R) transcripts, along with a modest elevation in D1 receptor (D1R) mRNA in the NAc. This D2R upregulation was associated with increased c-Fos activity in neurons expressing both D2R and D1R. Moreover, LP animals showed a higher proportion of D2R-positive neurons than D1R-positive neurons, hinting at a dominant role of the D2 pathway in LP offspring. The enhanced postsynaptic D2 receptor levels in the NAc could intensify motivation for palatable foods (Trifilief et al., 2013).

Additionally, we detected elevated TH and DAT transcript levels in the VTA, which implies higher dopaminergic neuronal activity or number. Prior findings support a correlation between TH mRNA expression and TH protein measured via immunohistochemistry (Paradis et al., 2017). This expanded dopaminergic drive from the VTA may potentiate medium spiny neuron (MSN) responses in the NAc, given the relevant DA receptor engagement.

Since the GABAergic signaling pathway was strongly impacted in the enrichment analysis, we next examined MSN electrophysiological features. MSNs make up approximately 95% of the striatal population (Kemps et al., 1971) and receive dopaminergic input from the VTA (Gerfen, 2010). Our focus was on the NAc shell, which is crucial for motivational processes, unlike the core subregion, linked to cognitive domains (Salgado and Kaplitt, 2014).

Our data reveal that, in childhood, MSNs from LP offspring displayed a higher spike threshold, with tendencies toward increased rheobase and extended slow afterhyperpolarization, suggesting reduced excitability. This decreased intrinsic excitability corresponded with lower amplitudes of GABAergic miniature inhibitory postsynaptic currents (mIPSCs), implying less effective GABA transmission between MSN or other striatal interneurons (Gerfen, 2010). Inadequate basal ganglia information processing may ensue if MSN excitability and GABA release are both altered. Furthermore, changes in MSN excitability, combined with heightened D2 receptor signaling, might explain why these pups displayed an elevated preference for palatable foods. However, our electrophysiological measurements did not distinguish between D1- and D2-positive MSNs. It is known that D1R-MSNs (direct pathway) and D2R-MSNs (indirect pathway) can be co-expressed in the NAc (Gangarossa et al., 2013), each influencing behavior differently (Falz et al., 2018; Wilson et al., 2017). In the dorsal striatum, D1R-MSNs are typically less excitable (i.e., require a higher rheobase) than D2R-MSNs (Gertler et al., 2008), although little is documented about NAc-specific properties. We observed a trend toward increased rheobase in P25 LP offspring, which could imply a larger D1R-MSN population in the probed region. Preliminary immunostaining (c-Fos analysis) in the NAc core and shell following a D1R agonist or D2R antagonist suggested more D1R and D2R cells in LP animals, aligning with the elevated receptor transcript levels at P25. Meanwhile, a decreased D1R/D2R ratio of c-Fos-positive cells points to fewer D1R-MSNs in the LP group, consistent with qPCR findings. Although these results warrant caution—given only partial immunoreactivity analyses—the overall picture indicates a broad remodeling in NAc MSN properties in response to perinatal malnutrition. Additional investigations are required to clarify the distinct roles of each pathway in palatable food consumption under nutrient restriction.

### 3.2. Impact of Prenatal LP Diet on Adolescence

During adolescence, both LP and CD animals initially exhibited similarly elevated preferences for palatable food over the first three days. However, after repeated exposure, LP rats progressively lost interest in the high-fat diet, thereby reducing their total caloric intake. To our knowledge, a diminished preference for palatable food specifically in adolescence has not been previously documented under maternal protein restriction. Nevertheless, our earlier work with maternal high-energy overnutrition mirrored this phenomenon (Paradis et al., 2017), suggesting that a waning hedonic response might underlie the reduced fat preference in LP animals. Because adolescence represents a period of robust neuronal remodeling (Spears, 2000), including changes in reward circuits (Naneix et al., 2016; Vendresculo et al., 2010), an unbalanced perinatal diet could disrupt the typical adolescent reorganization of hedonic food perception.

In parallel with behavioral data, FGSEA of RNA-seq highlighted dopaminergic pathway changes concentrated in the VTA. We confirmed increased DAT and D2R mRNA (with a trend toward heightened TH) by qPCR. By contrast, dopaminergic markers in the NAc largely normalized at this stage. However, the gene set enrichment pointed to continued perturbation of GABAergic pathways within the NAc in adolescent LP animals.

We then recorded electrophysiological parameters in MSNs at P50. In general, MSN features appeared nearly unchanged between groups, although we noticed a tendency toward a shorter action potential, implying a reduced postsynaptic transmission duration. Concomitantly, there was a significant elevation of miniature GABAergic current amplitudes, suggesting a mild increase in inhibitory tone impinging on MSNs. Previous work has shown that maternal malnutrition can lead to changes in neuronal excitability in other regions, such as the hippocampus, which displays fewer excitatory synapses and dendritic spines (Zhang et al., 2013).

Thus, LP animals demonstrated a faster decline in palatable food interest in adolescence, in contrast to the hyper-responsive behavior noted in early life. It appears that a normal chow diet provided in childhood may guard against exaggerated hedonic intake in adolescence. Possibly, an elevated dopaminergic input from the VTA (reflected by higher TH transcripts), alongside normalized DA receptor expression in the NAc and enhanced phasic GABA signaling in MSNs, underlies this reduced incentive for fatty foods. Our findings suggest that a three-week period of regular chow feeding post-weaning might “reprogram” reward circuits and confer resilience against acute palatable food challenges during adolescence.

### 3.3. Impact of Prenatal LP Diet on Adulthood

By adulthood, LP rats displayed an initial spike in their preference for palatable food on the first day of testing. However, their preference quickly normalized, matching that of CD rats by the second presentation. No significant hyperphagia was observed, despite the fact that LP animals still did not fully catch up in body weight. These results appear to contrast with prior reports of increased energy intake and higher palatable food preference in adult IUGR rats (Alves et al., 2015; Bellinger, 2004; Dalle Molle et al., 2015; Manuel-Apolinar et al., 2014; de Melo-Martimiano et al., 2015). In certain studies, dams were subjected to only one week of 50% caloric restriction in late gestation, whereas our design involved protein restriction throughout gestation and lactation. Such discrepancies highlight the significance of timing, duration, and the nature of nutritional restriction (Alves et al., 2015; Dalle Molle et al., 2015; Manuel-Apolinar et al., 2014) and suggest possible strain-specific effects (Bellinger, 2004; Melo-Martimiano et al., 2015). In one mouse model of perinatal protein restriction, for instance, adult offspring showed lower sugar preference (Vucetic et al., 2010), underscoring divergent outcomes based on species, strain, and experimental paradigms.

Several studies have documented alterations in dopamine-related gene expression caused by perinatal malnutrition (Alves et al., 2015; Dalle Molle et al., 2015; Laureano et al., 2016, 2018; Melo-Martimiano et al., 2015; Manuel-Apolinar et al., 2014; Vucetic et al., 2010). Our RNA-seq FGSEA revealed significant dopaminergic pathway modulation in the NAc, further supported by qPCR evidence of elevated D1R1a and unaltered D2R expression. This outcome aligns with Melo-Martimiano et al. (2015), who observed similar changes in DA receptor gene expression in the NAc of perinatally protein-restricted rats, but in their model, these changes correlated with pronounced food reward behaviors.

Our adult LP offspring also exhibited modified MSN electrophysiological properties in the NAc shell. We observed an increased rheobase and a more rapid rising phase of the action potential (with a slight trend for reduced spike duration), implying lower postsynaptic efficacy and decreased MSN excitability. Additionally, we detected a modest rise in the NMDA/AMPA receptor ratio, indicating shifts in synaptic plasticity mechanisms at excitatory synapses. With respect to GABAergic input, we noted no alterations in tonic GABA currents, but we did record enhanced miniature GABAergic current amplitudes, consistent with changes in GABAA receptor-mediated phasic inhibition. Interestingly, unlike in childhood, these GABAergic and excitatory synaptic adaptations during adulthood occurred alongside normalized food preferences. Thus, a notably different pattern of circuit remodeling emerges in adulthood: near-baseline dopaminergic activity in the VTA, elevated D1R expression in the NAc, reduced MSN excitability, and augmented phasic GABA currents—all converging to a state with typical eating behaviors.

Hence, despite evidence of lasting alterations in dopaminergic signaling and MSN physiology in the NAc attributable to perinatal malnutrition, adult offspring ultimately display eating preferences that are broadly similar to controls. This normalization likely arises from compensatory changes across multiple brain areas implicated in feeding regulation. Beyond the NAc and VTA, perinatal malnutrition can induce hypothalamic molecular alterations (Remmers, 2008; Vucetic et al., 2010), and these regions are extensively interconnected (Berthoud, 2002; Cassidy and Tong, 2019; O’Connor et al., 2015). Moreover, changes in electrophysiological traits have been documented in brain structures such as the dentate gyrus (Austin et al., 1986; Bronzino et al., 1997) and c-Fos activation shifts in the amygdala (Da Silva et al., 2016) and hippocampus (Zhang et al., 2010), potentially reshaping emotional responses to food and food-related memory. Together, these interrelated adjustments across numerous circuits may collectively maintain normal food intake if a standard diet is introduced post-weaning.

It is noteworthy that providing a standard diet after weaning, even following perinatal malnutrition, seems to trigger compensatory processes early on, particularly in offspring of LP dams. Although our study monitored palatable food intake every 24 hours, we cannot rule out subtler alterations in how animals experience “liking” versus “wanting.” Previous work has illustrated variations in meal patterns, satiety behaviors, and feeding frequency in other models of perinatal dietary imbalance (Breton et al., 2009; Coupé et al., 2012; Orozco-Solis et al., 2009; Wright et al., 2011).

Depending on the behavioral test employed, differences in motivation for rewarding foods have been reported (Romani-Pérez et al., 2017). Future studies are thus required to refine our understanding of how these circuits interact at a finer scale, mapping brain region cross-talk to specific facets of feeding motivation. This additional knowledge would deepen insights into the compensatory strategies by which normal eating behaviors are reestablished despite marked structural and functional neural changes incurred by perinatal undernutrition.

## 4. Conclusions

Our study aimed to investigate the developmental changes occurring in reward circuits in a perinatal protein restriction model, spanning from the weaning stage to adulthood. Through the use of comprehensive techniques, we have gained novel insights into the underlying mechanisms driving the evolution of palatable food preference observed in offspring exposed to protein restriction.

During childhood, we observed a positive correlation between an elevated preference for palatable food and increased dopaminergic activity in the ventral tegmental area (VTA). Additionally, gene expression analysis revealed higher levels of D2 receptors (D2R) and a relatively lower proportion of D1 receptors (D1R) in the nucleus accumbens (NAc). Moreover, a reduced excitability of medium spiny neurons (MSN) in the NAc, accompanied by a decrease in inhibitory postsynaptic current (IPSC) amplitude, was associated with the heightened preference for palatable food during this developmental stage.

In adolescence, a decline in the preference for palatable food was observed, suggesting the presence of protective mechanisms against the hedonic impact of such foods. This phenomenon was correlated with an increase in IPSC amplitude in NAc MSN and elevated dopaminergic activity in the VTA.

By adulthood, no significant difference in palatable food preference was observed; however, the analyzed parameters deviated from basal levels. Electrophysiological measurements revealed distinct functional characteristics in NAc MSN of offspring exposed to protein restriction, including reduced excitability, higher D1R gene expression, increased IPSC amplitude, and an elevated NMDA/AMPA ratio (markers of synaptic plasticity). These findings suggest the presence of compensatory mechanisms in other brain structures involved in the regulation of food consumption.

It is important to highlight that although preferences for fatty and sugary foods are transient during childhood, a balanced diet (chow diet) during early life leads to a decreased preference for such foods during adolescence and a return to normal levels in adulthood. Compensatory mechanisms are likely at work, allowing for adaptive behavior. However, further investigations are required to explore the long-term functioning of this “modified” circuit and its response to stress.

## 5. Materials and Methods

### 5.1. Ethics Statement

All experiments were performed in accordance with the guidelines of the local welfare committee, the EU (directive 2010/3/EU), the Institut National de la RechercheAgronomique (Paris, France), and the French Veterinary Department (A44276). The protocol was approved by the institutional ethic committee and registered under reference APAFIS 7953. Every precaution was taken to minimize stress and the number of animals used in each series of experiments.

### 5.2. Animals and Diets

The animals were housed in a reverse light cycle with a day cycle of 12h00 starting at 19h00 until 7h00 of the following day. The temperature of the housing room was controlled and maintained around 22±2°C with access to food and water ad libitum. Female Sprague Dawley rats (body weight: 240-290 g on day 1 of gestation (G1) were purchased directly from Janvier (Le Genest Saint Isle, France). They were housed in individual cages from arrival and fed either a control diet (CD) (20% protein) or a low protein diet (LP) (8% protein) (isocaloric diets) (INRAE, UPAE, Jouy en-Josas, France) during gestation and lactation (see Table 1: Diet compositions). At birth, litter size was adjusted to eight pups per dam with a 1:1 male:female ratio (Figure 1 B). From weaning (P21), pups born to CD and LP dams were fed a standard diet until the end of the experiment (see Table 1). The body weight of the pups was monitored regularly throughout the experiment. In this work we present the data on the male and female offspring.

**TABLE 1:** Diet compositions.

| Nutriments | Control Diet (Ctrl) | Low Protein Diet (LP) | Standard Diet (SD) | Western Diet (WD) |
| --- | --- | --- | --- | --- |
| Cellulose | 5 | 5 | 3.9 | 5 |
| Fat | 7 | 7 | 3.1 | 21.0 |
| Sucrose | 10 | 11.9 | 2.0 | 29.45 |
| Casein | 20 | 8 | 16.1 | 22.0 |
| Dextrose | 13.2 | 15.7 | 12.2 | 10.0 |
| Starch | 39.7 | 47.3 | 45.8 | 5.0 |
| Total energy, kcal/100g | 352.7 | 352.2 | 301.4 | 466.3 |

**TABLE 2.** Passive and active membrane properties.

|  | Childhood (PN18-PN28) |  | Adolescence (PN48-PN62) |  | Adulthood (PN89-PN106) |  |
| --- | --- | --- | --- | --- | --- | --- |
|  | Control (n=30) | IUGR (n=26) | Control (n=16) | IUGR (n=20) | Control (n=12) | IUGR (n=17) |
| Cm (pF) | 12.96 (± 0.130) | 13.21 (± 0.18) | 12.946 (± 0.24) | 12.97 (± 0.23) | 12.58 (± 0.23) | 12.53 (± 0.62) |
| Tau CM (μs) | 1.20 (± 0.04) | 1.24 (± 0.05) | 1.32 (± 0.07) | 1.41 (± 0.07) | 1.26 (± 0.08) | 1.36 (± 0.10) |
| Input Resistance (mOhm) | 185.9 (± 12.12) | 192.3 (± 12.47) | 112.6 (± 5.97) | 123.6 (± 8.22) | 163.9 (± 12.40) | 138.2 (± 15.29)(*) |
| Resting Membrane Potential (mV) | -79.10 (± 1.16) | -78.56 (± 1.13) | -81.20 (± 1.15) | -79.10 (± 1.56) | -78.72 (± 1.15) | -78.87 (± 1.63) |
| Rheobase (pA) | 91.33 (± 6.79) | 109.23 (± 6.72) | 172.50 (± 13.56) | 151.00 (± 12.83) | 100.83 (± 6.09) | 141.18 (± 13.39)** |
| Spike threshold (mV) | -39.65 (± 0.53) | -37.05 (± 0.50)** | -38.34 (± 1.07) | -39.03 (± 0.92) | -39.31 (± 1.12) | -38.93 (± 0.47) |
| Delay for first spike (ms) | 372.66 (± 13.12) | 354.11 (± 16.35) | 379.75 (± 19.10) | 384.78 (± 19.41) | 392.25 (± 30.94) | 379.81 (± 12.43) |
| Spike amplitude (mV) | 81.28 (± 0.91) | 80.18 (± 0.91) | 85.91 (± 1.28) | 86.94 (± 1.11) | 83.36 (± 1.90) | 85.73 (± 1.05) |
| Spike duration (ms) | 3.19 (± 0.14) | 3.00 (± 0.10) | 3.17 (± 0.18) | 2.75 (± 0.10) (*) | 3.31 (± 0.15) | 2.88 (± 0.16)(*) |
| Spike rise time (ms) | 0.73 (± 0.04) | 0.73 (± 0.03) | 0.73 (± 0.08) | 0.67 (± 0.03) | 0.86 (± 0.05) | 0.68 (± 0.04)** |
| Spike decay time (ms) | 2.45 (± 0.11) | 2.27 (± 0.08) | 2.44 (± 0.16) | 2.08 (± 0.09) (*) | 2.45 (± 0.12) | 2.19 (± 0.13) |
| Ratio spike rise/decay time | 0.30 (± 0.01) | 0.33 (± 0.02) | 0.33 (± 0.06) | 0.33 (± 0.02) * | 0.35 (± 0.02) | 0.32 (± 0.02) |
| fAHP amplitude (mV) | 10.24 (± 0.35) | 10.07 (± 0.27) | 9.62 (± 0.35) | 10.15 (± 0.36) | 10.53 (± 0.65) | 10.07 (± 0.36) |
| fAHP duration (ms) | 13.82 (± 1.19) | 12.23 (± 1.28) | 16.41 (± 1.98) | 14.74 (± 1.86) | 11.15 (± 2.74) | 13.58 (± 1.97) |
| fAHP dV/dt (mV/ms) | 1.11 (± 0.20) | 1.43 (± 0.27) | 0.94 (± 0.28) | 2.05 (± 0.68) | 2.31 (± 0.59) | 1.85 (± 0.64) |
| sAHP duration (ms) | 60.84 (± 4.64) | 73.99 (± 5.28)* | 66.62 (± 4.73) | 68.75 (± 3.81) | 80.64 (± 5.00) | 67.32 (± 3.56) * |
| AP frequency (Hz) (rheobase +30 pA) | 10.93 (± 0.55) | 9.85 (± 0.50) | 8.88 (± 0.58) | 9.20 (± 0.44) | 8.67 (± 0.45) | 9.06 (± 0.60) |

### 5.3. Behavior (Free Choice Test for Food)

Three critical developmental periods were studied (P21 to P25: juvenile, P43 to P50: adolescence, and P93 to P100: young adult) (Figure 1 A). For each period, 24 male pups (n=12 per group) were randomly selected and placed in an individual cage to perform a free-choice test with highly palatable food (see Table 1). Different pups were used for each period (Figure 1B). This test was used to specifically study the attraction towards palatable food. After one day of habituation to the presence of food on both sides of the cage, the test was performed over 3 days at P22, and over 6 days at P44 and P94 (Figure 1 A). In detail, different pups were housed individually for 4 days at P21 and 7 days at P43 and P93: day 1, habituation phase with a standard diet on both sides of the cage; day 2 to day 4 or day 7, rats were given a free choice between a standard diet and a Western diet (ABdietWoerden, The Netherlands) (see Table 1). The position of the two different foods was switched daily to avoid positional preference bias. The consumption of the different foods was recorded daily at 11:00 am. The fat preference score was calculated as the ratio of the amount of “palatable food” consumed to the total amount of food consumed in 24 hours.

### 5.4. Tissues Collection and Blood Sampling

The day after the last day of the free choice test (P25, P50 or P100), half of the rats (n=6 per group) were rapidly euthanized between 9:00 and 12:00 am by CO2 inhalation. Blood was collected in tubes with EDTA (LaboratoiresLéo SA, St Quentin en Yvelines, France) and centrifuged at 2.500 g for 15 min at 4°C. Plasma was frozen at −20°C. The brain was rapidly removed and placed in a brain matrix (WPI, Sarasota, FL, USA rat 300-600g). First the hypothalamus was dissected (according to Paxinos’ atlas coordinates: −1.0 to −4.5 mm from Bregma) then, for each rat, two coronal slices of 2 mm thickness at the level of Nucleus Accumbens and another one at the level of the Ventral Tegmental Areawere obtained. Samples of the right and left NAc and VTA (four samples in total per animal) were rapidly obtained using two different biopsy punches (Stiefel Laboratories, Nanterre, France) (diameter of 4 mm for the NAc and 3 mm for the VTA). The samples were snapped frozen in liquid nitrogen and stored at −80°C for subsequent determination of gene expression by DGE – RNA seq and TaqMan (as already described in Paradis et al. 2017).

As previously described in Paradis et al., 2017; plasma samples collected from P25, P45 and P95 rats were used to measure plasma glucose, NEFA (non-esterified fatty acids), insulin and leptin. Glucose and NEFA were measured by colorimetric enzymatic reactions with specific kits (glucose and NEFA PAP 150 kits, BioMérieux, Marcy l’Etoile, France). Hormones were assayed with specific ELISA kits following the manufacturer’s instructions for insulin and leptin (rat / mouse insulin ELISA kit, rat leptin ELISA kit, Linco Research, St. Charles, MO).

### 5.5. Gene Expression Analysis

- DGE RNA seq: According to the preference tests, this analysis was performed only on male samples. Analysis of RNA-seq and DGE-seq were performed on sample of VTA and NAc following the same protocol as already described in Kilens et al. 2018. RNA was isolated from snap-frozen NAc and VTA-enriched samples using the NucleoSpin RNA/protein kit (Macherey-Nagel, Hoerdt, France). Total RNA was submitted to DNase digestion following the manufacturer’s instructions, the quantity was estimated by the 260/280 nm UV absorbance, and the quality was assessed using the Agilent 2100 Bioanalyzer System, the RNA integrity number (RIN) was then calculated. Samples with a RIN below 7 were discarded. The libraries were prepared from 10 ng of total RNA. The mRNA poly(A) tails were tagged with universal adapters, well-specific barcodes and unique molecular identifiers (UMIs) during template-switching reverse transcriptase. Barcoded cDNAs from multiple samples were then pooled, amplified and tagmented using a transposon-fragmentation approach which enriches for 3′ends of cDNA. A library of 300–800 bp was run on an Illumina HiSeq 2500 using a Hiseq Rapid SBS Kit v2-50 cycles (ref FC-402-4022) and a Hiseq Rapid PE Cluster Kit v2 (ref PE-402-4002). Read pairs used for analysis matched the following criteria: all 16 bases of the first read had quality scores of at least 10 and the first 6 bases correspond exactly to a designed well-specific barcode. The second reads were aligned to RefSeq2. A gene set functional enrichment analysis (FGSEA) method was used to identify differentially regulated pathways. Enrichment was performed using the Gene Ontology (GO) [ref the gene ontology 2019], Kyoto Encyclopedia of Genes and Genomes (KEGG) [Kanehisa and Goto 2000] and Reactome [Jassal et al., 2020] databases. Pathways with an FDR < 0.01 were retained for further analysis.
- Gene expression by Real-time Quantitative Polymerase Chain reaction (RT-qPCR) This analysis was performed only on male samples. The exact same samples were used for RT-qPCR. One microgram of total RNA was reverse transcribed into cDNA using High capacity RT kit (Applied Biosystems, Foster City, CA, USA) in a total volume of 10 µl. PCR was carried out using Bio-Rad iCycler iQ system using the qPCR SYBR Green MasterMix (Fermentas, Courtaboeuf, France). Quantitative PCR consisted of 45 cycles, 30 s at 95°C and 30 s at 60°C each.. The assays per sample were n = 6 (n = 5 for CD group at P25).

### 5.6. Electrophysiology

This analysis was performed only on male in separate group of animals (Figure 1A), electrophysiological properties of Medium Spiny Neurons (MSN) were studied in the nucleus accumbens (shell part) (Figure 4). Experiences were carried out on pups of both groups for the 3 critical developmental periods P18 to P28: juvenile; PN48 to P62 adolescence and P89 to P106 young adult (Figure 1A). Pups did not do any behavioral test before electrophysiology study (Figure 1A).

- Tissue preparation Rats were anesthetized with sub-lethal dose of pentobarbital and killed by decapitation. Brains were rapidly removed and sectioned into 300 µm-thick slices in the sagittal plane with a Leica VT 1000S microtome (LEICA Microsystem, Germany) in an ice-cold extra-cellular solution of the following composition (in mM): 25 glucose, 125 NaCL, 2.5 KCl, 2 CaCl2, 1 MgCl2, 25 NaHCO3, 1.25 NaH2PO4 (gassed with 95 % O2/ 5% CO2) (about 295 mosmol). Slices were then left for equilibration for ∼ 1 hour in a bath at 34 °C in the same extra-cellular solution (gases with 95% O2/ 5% CO2).
- Whole cell recordings Single slices were transferred to a recording chamber and perfused continuously (pump minipuls 3, GILSON INC, Middleton, WI, USA) with extra cellular solution (see above) at room temperature and saturated with 95% O2/5%CO2. Neurons were visualized with Olympus microscope (Olympus BX51WIF, Olympus corporation, Japan) and a X40 water-immersion objective (Olympus U-CAMD3, Olympus corporation, Japan). Whole cell recordings were performed using borosilicate pipettes (Glass capillaries GC150TF-10, Harvard apparatus, USA) prepared with a micropipette puller (PC-10, Narishige Co, LTD, Japan). Electrode (7-8Mohm) were filled with a solution containing the following (in mM): 122Kgluconate, 13 KCl, 0.3 EGTA, 10 HEPES, 4 MgATP, 0.3 NaGTP, 10 phosphocreatine, Adjusted to pH 7.35 and 298 mosmol for protocol of action potential and NMDA/AMPA ratio recordings or with a CsCl solution containing the following (in mM): 135 CsCl, 0.3 EGTA, 10 HEPES, 4 MgATP, 0.3 NaGTP, 10 Phosphocreatine in the case of minis recordings. MSN neurons were distinguished from other neurons (cholinergic and interneurons) based on their electrophysiological properties. Protocol of action potential of all neurons recorded was assessed at the beginning of every experiment. Data were recorded using a HEKA amplifier and software (HEKA EPC10 USB double, HEKA Instrument Inc, New-York, USA). Signals were digitized at 1 kHz. Recordings with KGluconate-filled and CsCl –filled pipettes were corrected for a junction potential +18 mV and −7 mV respectively. Series resistance was monitored throughout the experiment by a brief voltage step of −10 mV at the end of each recording. Data were discarded when the series resistance increased or decreased by >20%.
- Drugs Unless otherwise stated, all drugs used were purchase from Tocris and prepared as concentrated stock solution and stored at −20°C. On the day of experiment, drugs were diluted and applied through the bath perfusion system. GABAergic synaptic transmission was blocked for minis recording. GABAA receptors were blocked with 100 µM picrotoxin (PTX) dissolved in extracellular solution. Stock PTX was dissolved in ethanol and then added in the external solution at a final ethanol concentration of 0.01–0.02%. NMDA or AMPA/Kainate receptors were blocked with 50 µM D-AP5 (C5H12NO5P) and 10 µm CNQX (C9H4N4O4) solution respectively.
- Action potential recording and EPCSs recording: Active membrane properties The current was clamped to membrane potential (0 pA injected). Progressive current injections were done during 500 ms at 0.1Hz (beginning of the protocol with injection of −150 pA current and then progressive current injection with steps of 10 pA to firing neuron). Action potential parameters recordings were analyzed following a current injection of +30 pA above rheobasis: membrane potential, amplitude of action potential, rise and decay time duration, amplitude of hyperpolarisation, duration to reach hyperpolarisation amplitude, frequency of action potential (as already described in Paillé et al., 2007).
- NMDAR-AMPAR ratio For the NMDAR/AMPAR ratio experiments, MSNs were voltage clamped at −80mV and +40mV to record, respectively, AMPAR mediated and NMDAR-mediated EPSCs. The NMDAR EPSCs component was individuated by using the kinetic method, considering the peak amplitude at 50 ms after the beginning of the event. The NMDAR/AMPAR ratio was calculated by dividing the NMDAR peak by the AMPAR peak. Presynaptic cells were stimulated by a bipolar stimulating electrode placed in the cortex near to the recording electrode in the nucleus accumbens (approximatively 100 µm). AMPAR and NMADAR EPCS peak amplitudes were measured from the average of 10 traces.
- Minis analysis spontaneous EPSCs Phasic and Tonic GABA current Whole cell patch clamp recording were made in distinct slice than the slices used for AP and NMDA/AMPA ratios. Spontaneous phasic and GABAAR-mediated current measurement was made as previously described by Valtcheva et al., 2017 (Valtcheva et al., 2017). Briefly, spontaneous phasic and tonic GABAAR-mediated currents were measured using a Cs+-high-chloride based intracellular solution. Phasic and tonic GABAergic components were estimated after inhibition of ionotropic glutamatergic receptors by adding D-AP5 (50 µM) and CNQX (10 µM) at the beginning of the experiment. Phasic and tonic components were analyzed during at least 3 consecutive 25 s recording segments before and after pharmacological treatments. Concerning phasic GABAAR-mediated current, spontaneous IPSCs (sIPSCs) were identified using a semi-automated amplitude threshold-based detection software (Mini Analysis 6.0.7 Program, Synaptosoft, Fort Lee, NJ, USA) and were visually confirmed. Concerning tonic GABAAR-mediated current, the holding current was measured based on an average of 4 recording segments preceding drug application. After pharmacological treatment, we determined a new Ihold and ΔIhold corresponded to the tonic component affected by the drug. Picrotoxin was added in the bath at the end of the recording to estimate the magnitude of the tonic GABAergic signaling.

### 5.7. D1/D2 Neurons Quantification

- Drugs SKF81297 (5.0 mg/kg), Raclopride (0.3 mg/kg) were purchased from Tocris and dissolved in 0.09% NaCl (saline solution). Drugs were administrated with intra peritoneal injections on day 4 after a free choice test with high fat food on P25 pups (same test as already described above).
- Tissue collection and preparation In a separate group of animals (Figure 1A), 24 males (n=12 per group i.e 12 CD pups and 12 LP pups) were randomly selected and assigned to a specific drug injection Raclopride, SKF81297 or vehicle saline solution (n=4 for each subgroup). P25 rats received intraperitoneal injection (0.2ml/100g). One hour and a half after drug injection, rats were anesthetized (after buprenorphine injection 15min before the sacrifice) with isoflurane and perfused with a transcardial physiological saline perfusion followed by ice-cold 4% paraformaldehyde in phosphate buffer (PB, pH 7.4). The brains were rapidly removed, immersed in the same fixative for 1 day at 4°C, and finally stored in 30% PB sucrose for 24-48h. The brains were then frozen in isopentane at −60°C, and finally stored at −80°C until use. The NAc was cut into 40 µm free floating serial coronal sections with a cryostat (Microm, Microtech, Francheville, France). Free-floating slices were stored at −20°C in an antifreeze mixture (glycerol 25%, ethylene glycol 25% and PBS 0.2M 50%) until their utilization.
- Immunohistochemistry c-Fos Wells containing serial NAcfree-floating sections were first well rinsed in PB 0.1M to eliminate anti-freeze solution. Then endogenous peroxidase was deactivated by incubating slices in peroxidase block (0.3% H2O2 solution) for 30 min with gentle agitation. Slices were rinsed in PB 0.1M and then incubated 2h in blocking buffer (PB 0.1M/3% DNS/0.25%Triton X-100). Free-Floating slices were incubated over week-end at 4°C with a mixture of the following primary antibody: anti-cfos (1:10000; Santacruz sc-52) in blocking buffer (PB 0.1M/3% DNS/0.25%Triton X-100). After incubation with primary antibody and subsequent washing in PB 0.1M, free-floating slices were incubated with second antibody: goat anti rabbit biotin (1:1000; Life Tech A24541) for 2h. Sections were rinsed and then incubated with ABC kit (Eurobio Vector PK-7100) for 30 min. After rinsing, sections were placed in solution for DAB reaction (Eurobio Vector SK-4100) with Nickel. Reaction once the background is high enough was stopped by placing sections into water and methyl green colorations were done. Then sections were air-dried during 2 days and mounted in dehydrated medium.
- D1/D2 neurons count in Nucleus Accumbens: For each rat, c-Fos-positive cells were counted at three different rostrocaudal levels of the NAc (distance relative to Bregma: + 0.84 mm, +1mm and +1.16 mm referring the Paxinos and Watson atlas). For the left and the right side, a digitized picture comprising the whole Striatum and NAc were obtained using X20 magnification of a NanoZoomer-Xr Digital slide scanner C12000 (Hamamatsu, Japan). To obtain the least biased estimate of the total number of neurons, we used the dissector principle and random systematic sampling (Sterio, 1984; Coggeshall, 1992). First, a line was drawn around the core and the shell area of NAc for each section. Using the NIH Image J software (cell counter plugin), C-Fos immunoreactive body cells were counted based on a stereological technique (unbiased quantization technique) to limit the bias of “over-counting”. A counting grid was randomly positioned on the image (“systematic random”), each small square having a length of 100µm and only the c-Fos positive cells present in one square out of four were counted by an observer blind to the rat number (Figure 7). Once the counts done, the total density of the area was extrapolated according to the area of the analyzed structure. Data are expressed as mean of labeled cells/mm² ± SEM.

### 5.8. Statistical Analysis

Results are expressed as mean ± SEM in figures and tables. Mann-Whitney non-parametric tests were done for the analysis of body weight, the analysis of fat preferences and the analysis of electrophysiological parameters at different time points. For the plasma sample analysis, a non-parametric Mann and Whitney test were also performed. These statistical analyses were performed using Prism 9.0 software (GraphPadSofware Inc., La Jolla, CA, USA).

For RNA-seq analysis, statistical analyses were performed using the software Prism (Graphpad) or R. Statistical test were done as already describe in Kilens et al. 2018. DESeq2 was used for analysis considering RNA-seq data were following a negative binomial distribution. Other statistical tests were performed considering their specific assumption and hypothesis, notably for Pearson correlation’s test and homogeneity χ2 tests.

## Author Contributions

JP and CG conducted the experiments and took part in both the discussion and manuscript writing. MF, VA and DM performed the DGE-RNAseq and contributed to the discussion and writing. AP and PdC helped design the experiment and participated in the discussions. PP also contributed to the experimental design, took part in the discussions, and wrote the manuscript. VP conceived and performed the experiments, analyzed the data, and wrote the manuscript.

## Funding

Pays de la Loire region funding PARIMAD (VP). Nantes University PhD salary (JP).

## Acknowledgments

We acknowledge the IBISA MicroPICell facility (Biogenouest), member of the national infrastructure France-Bioimaging supported by the French national research agency (ANR-10-INBS-04).

## Conflicts of Interest

The authors declare no conflict of interest.

## Notes

### Competing Interest Statement

The authors have declared no competing interest.

